# SynToxProfiler: an approach for top drug combination selection based on integrated profiling of synergy, toxicity and efficacy

**DOI:** 10.1101/693010

**Authors:** Aleksandr Ianevski, Alexander Kononov, Sanna Timonen, Tero Aittokallio, Anil K Giri

## Abstract

Drug combinations are becoming a standard treatment of many complex diseases due to their capability to overcome resistance to monotherapy. Currently, in the preclinical drug combination screening, the top hits for further study are often selected based on synergy alone, without considering the combination efficacy and toxicity effects, even though these are critical determinants for the clinical success of a therapy. To promote the prioritization of drug combinations based on integrated analysis of synergy, efficacy and toxicity profiles, we implemented a web-based open-source tool, SynToxProfiler (Synergy-Toxicity-Profiler). When applied to 20 anti-cancer drug combinations tested both in healthy control and T-cell prolymphocytic leukemia (T-PLL) patient cells, as well as to 77 anti-viral drug pairs tested on Huh7 liver cell line with and without Ebola virus infection, SynToxProfiler was shown to prioritize synergistic drug pairs with higher selective efficacy (difference between efficacy and toxicity level) as top hits, which offers improved likelihood for clinical success.

## Introduction

High throughput screening (HTS) of approved and investigational agents in preclinical model systems has been established as an efficient technique to identify candidate drug combinations to be further developed as safe and effective treatment options for many diseases, such as HIV, tuberculosis and various types of cancers [1, 2]. Currently, the selection of top combinations for further development often relies merely on the observed synergy between drugs, while neglecting their actual efficacy and potential toxic effects, that are the other key determinants for the therapeutic success of drugs in the clinics [3]. Notably, around 20% of drugs fail in the early development phase because of safety concerns (non-tolerated toxicity), and over 50% fail due to lack of sufficient efficacy [4]. Further, a recent study argued that many clinically-used anticancer combination therapies confer benefit simply due to patient-to-patient variability, not because of drug additivity or synergy [3], indicating that even non-synergistic combinations may be beneficial for therapeutic purposes if they have a high enough efficacy and low enough toxicity profiles. To make a better use of these various components of drug combination performance already in preclinical HTS experiments, we implemented, to the best of our knowledge, the first web-tool, SynToxProfiler, which enables users to profile synergy, toxicity and efficacy of drug combinations simultaneously for the top hit prioritization and is also extendible for multi-drug (3 or more drugs) combination screening.

## Methods

### SynToxProfiler workflow

The SynToxProfiler web-application is freely available at https://syntoxprofiler.fimm.fi, together with example drug combination data, video tutorial and user instructions. SynToxProfiler enables ranking of drug combinations based on integrated efficacy, synergy and toxicity profiles (Fig.1). Therefore, for each drug combination, SynToxProfiler first calculates a normalized volume under dose-response surface to quantify combination efficacy based on dose–response measurements on diseased cells, e.g. patient derived primary cells (see Suppl. Fig. 1). Then, the combination synergy between each drug pair is estimated using one of the synergy scoring models: Highest Single-Agent [5], Bliss independence [6], Loewe additivity [7], or Zero Interaction Potency [8], as implemented in the SynergyFinder R-package [9]. Normalized volume under the dose-synergy surface is utilized to quantify final combination synergy score (Suppl. Fig. 1A). Next, using the measurements on control cells, if available, the normalized volume under dose–response matrix is calculated to estimate combination toxicity (Suppl. Fig. 1). Finally, SynToxProfiler ranks the drug combinations based on integrated combination synergy, efficacy and toxicity (STE) score. Alternatively, if measurement on control cells are not available, then the ranking of drug pairs can also be done based merely on combination synergy and efficacy. As a result of the interactive analysis, SynToxProfiler provides a web-based exportable report, which allows users to interactively explore their results (Fig. 1 and Suppl. Fig 2). An interactive example of web-based report is given at https://syntoxprofiler.fimm.fi/example. A more detailed description of the calculations and workflow is provided in the technical documentation, https://syntoxprofiler.fimm.fi/howto.

**Fig. 1.**
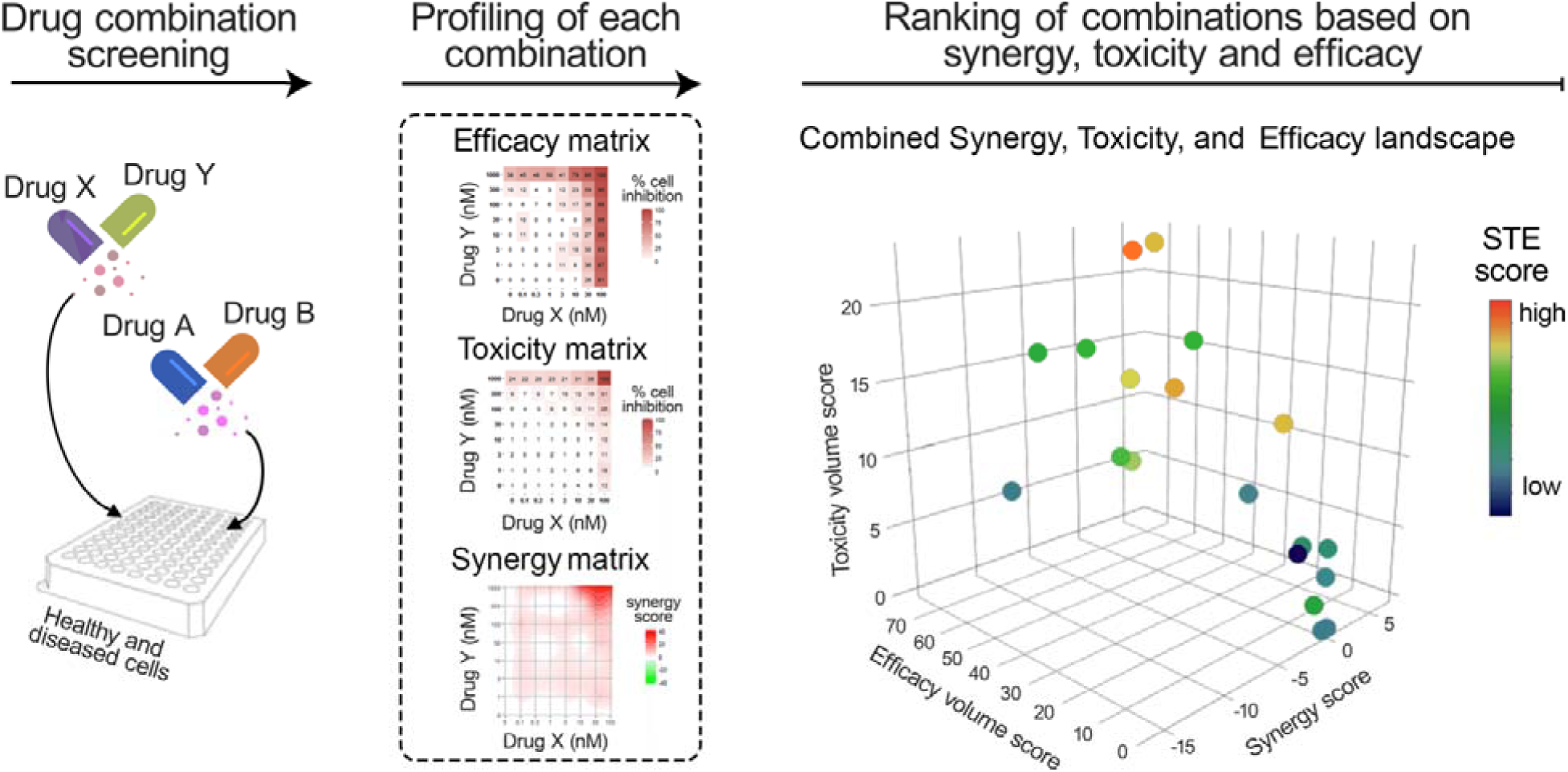
A schematic overview of SynToxProfiler. The dose-response data from drug combination screening, measured in both diseased (e.g. patient-derived cells) and healthy control cells (e.g. PBMCs), is provided as input to SynToxProfiler (left panel). Then, SynToxProfiler quantifies drug combination efficacy and synergy (using combination responses in diseased cells) as well as toxicity (using combination responses in control cells) for each drug pair (middle panel), and summarizes them into integrated synergy, toxicity and efficacy (STE) score. The STE score is further used to rank and visualize the drug pairs in 2D or 3D interactive plots (right panel).

### Calculation of normalized volume

The normalized volume under the dose-response surface is calculated while quantifying combination efficacy and toxicity based on measurements on diseased and control cells, respectively (Suppl. Fig. 1). Synergy score was calculated based on measurements on diseased cells as normalized volume under synergy matrix (excess matrix of combination responses over expected responses determined by one of the synergy models, such as Bliss). For each combination AB of drugs, A and B, the normalized volume under the dose-response surface V_AB_ is calculated as:

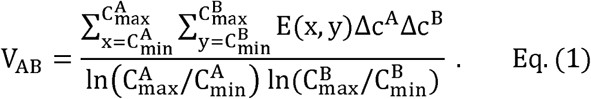

Here, c^A^_min_ and c^A^_max_ are the minimum and maximum tested concentrations of drug A, respectively, and c^B^_min_, and c^B^_max_ are those of drug B; Δc^A^ and Δc^B^ are the logarithmic increase in concentration of drug A and drug B between two consecutive measurements of dose-response matrix; and E(x, y) is the efficacy or toxicity levels at concentration x of drug A and at concentration y of drug B. The current approach for volume-based scoring normalizes for the different dose-ranges measured in different drug combinations, as commonly occurring in HTS settings. The extension of formulation for volume - based scoring of synergy, efficacy and toxicity profiles for multi-drug combinations (3 or more drugs) is given in the supplementary file.

### Ranking of drug combinations

SynToxProfiler ranks the drug combinations based on an integrative analysis of synergy, toxicity and efficacy, quantified as STE score. First, the difference in efficacy (E_AB_) and toxicity volume scores (T_AB_) is calculated for each drug combination to quantify a selective response in diseased cells, relative to that of control cells. We defined this difference as a selective efficacy score (sE_AB_) of a drug combination. This theoretical concept for selective efficacy has been adopted from the single drug dose-response assays, where the difference in normalized areas under the curve (AUC) between diseased and healthy cells is often used to calculate the patient-specific drug efficacies [10, 11]. The final STE score is given by averaging two different ranks of (i) combination synergy score (S_AB)_ (the higher is the synergy, the higher is the rank), and (ii) selective combination efficacy (sE_AB_) (the higher is selective efficacy, the higher is the rank):

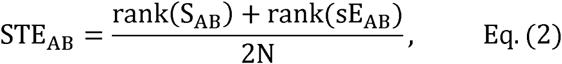

where S_AB_ and sE_AB_ are the synergy and selective efficacy scores, respectively, for a combination of drug A and B, calculated using the normalized volume under the dose-response surface; and N is the total number of drug combinations being tested. However, since calculation of STE score using the whole dose-response matrix may miss some of the top hit drug combinations with a narrow synergistic dose window, SynToxProfiler also offers the users a possibility to rank combinations based on the selective efficacy and synergy scores calculated only at the most synergistic area of the drug combination matrix (defined as the 3×3 concentration window with the highest synergy in the dose-response matrix), instead of the default full matrix calculation.

### Data submission and reporting

The default input of SynToxProfiler is a text or xlsx file that comprises annotations of each drug combination dose–response matrix, including drug names, concentrations, cell types (e.g. sample or control), and phenotypic responses (e.g. relative inhibition). The number of drug combinations provided in the input file is unrestricted. More information on the input file format is given in the website documentation (https://syntoxprofiler.fimm.fi/howto/). As the result, SynToxProfiler provides an interactive visualization of STE scores using bar charts, as well as 2- and 3-dimensional scatter plots. Publication-quality figures (e.g. heatmap for dose-response and synergy matrix, 2D and 3D scatter plot for different scores) can be exported in PDF files, as well as all the calculated scores can be downloaded in an xlsx file.

### Drug combination assay

The in-house drug combination testing was carried out at Institute for Molecular Medicine Finland (FIMM), in peripheral blood mononuclear cells (PBMCs) of a patient with T-cell prolymphocytic leukemia (T-PLL) and a healthy volunteer were used in accordance with the regulations of Finnish Hematological Registry and biobank (FHRB). The written informed consents were obtained from both participants and the study was carried in accordance with the principles of Helsinki declarations. Twenty combinations of drugs with different mechanisms of actions (see Supplementary Table S1) were tested on the PBMCs in 8×8 dose-response matrix assay as described previously [12, 13]. Briefly, 20 microliters of cell suspension along with compounds (in 8 different concentrations including zero dose) and their combinations were plated on clear bottom 384-well plates (Corning #3712), using an Echo 550 Liquid Handler (Labcyte). The concentration ranges were selected for each compound separately to investigate the full dynamic range of dose-response relationships. After 72 hours incubation at 37°C and 5% CO2, cell viability of each well was measured using the CellTiter-Glo luminescent assay (Promega) and a Pherastar FS (BMG Labtech) plate reader. As positive (total killing) and negative (non-effective) controls, we used 100 μM benzethonium chloride and 0.1 % dimethyl sulfoxide (DMSO), respectively, for calculating the relative efficacy (% inhibition).

The published dataset of 78 antiviral drug combinations was tested at the Integrated Research Facility, National Institutes of Allergy and Infectious Diseases (NIAID), in the Huh7 liver cells infected with Makona isolate, Ebola virus/H.sapiens-tc/GIN/14/WPG-C05, as described in the original study [14] (data available at https://matrix.ncats.nih.gov/matrix-client/rest/matrix/blocks/6323/table and https://matrix.ncats.nih.gov/matrix-client/rest/matrix/blocks/6324/table). Briefly, drugs in 50-µL of Dulbecco’s modified Eagle’s medium were transferred to the Huh7 cells seeded in black, clear-bottomed, 96-well plates 1 hour prior to inoculation with EBOV/Mak. After 48 hours of viral inoculation, drug combination efficacy was measured in triplicates with a 6 × 6 dose-response matrix design using CellTiter-Glo assays (Promega). The EBOV/Mak virus was detected using mouse antiEBOV VP40 antibody. For the toxicity measurements, the same CellTiter-Glo assay was performed on non–virus-infected Huh7 cells with 3 replicates for each drug concentration, and the assay was repeated at least twice for confirmation. We utilized 77 out of the 78 combinations for the present analysis, as colchicine-colchicine pair was removed because the inhibition levels were 100% for all the tested concentrations for the drug combination.

## Results

### SynToxProfiler prioritizes clinically useful drug combination as top hits for T-PLL cancer patient and Ebola virus infection

To demonstrate the performance of SynToxProfiler in prioritizing therapeutically-relevant synergistic combinations, we applied it to in-house drug screening data involving 20 drug combinations tested in one control and one T-PLL patient-derived cells. The T-PLL case study revealed that ranking of combinations based on the STE score successfully prioritizes both effective and safe drug pairs. For example, Cytarabine-Daunorubicin pair was identified as the top hit out of the tested combinations (Table 1, Additional File 1); this combination is widely used as approved induction therapy for acute myeloid leukemia treatment [15,16]. Ibrutinib-Navitoclax was ranked as the third-best combination for further study; this combination has shown promising results in phase II clinic trail (NCT02756897) for chronic lymphocytic leukemia (CLL), and recently suggested as first-line treatment for CLL [17].

**Table 1:** Ranking of 20 in-house measured combinations based on STE scores calculated from the most synergistic area of dose-response matrix in T-PLL and healthy control cells

| Drug1 (concentration range in nM) | Drug2 (concentration range in nM) | Synergy score ($S_{AB}$) | Efficacy score ($E_{AB}$) | Toxicity score ($T_{AB}$) | Selective efficacy score ($E_{AB}$) | STE score |
| --- | --- | --- | --- | --- | --- | --- |
| Cytarabine (0 -100) | Daunorubicin (0 - 1000) | 6.701 | 58.727 | 20.864 | 37.863 | 0.825 |
| Trametinib (0 -100) | S-63845 (0 - 25) | 4.202 | 48.4 | 12.427 | 35.973 | 0.825 |
| Ibrutinib (0 -1000) | Navitoclax (0 - 100) | 1.529 | 71.686 | 19.922 | 51.764 | 0.825 |
| Quizartinib (0 -100) | S-63845 (0 - 100) | 0.69 | 56.379 | 27.43 | 28.949 | 0.75 |
| Omacetaxine (0 -1000) | Ipatasertib (0 - 1000) | 0.188 | 70.096 | 11.361 | 58.735 | 0.725 |
| Gefitinib (0 -1000) | Omacetaxine (0 - 1000) | 0.693 | 69.355 | 17.071 | 52.284 | 0.725 |
| Clofarabine (0 -1000) | Idarubicin (0 -100) | 2.075 | 73.876 | 27.528 | 46.348 | 0.725 |
| Omacetaxine (0 -1000) | Alpelisib (0 - 1000) | -0.723 | 70.87 | 10.947 | 59.923 | 0.675 |
| Clofarabine (0 -1000) | Prexasertib (0 - 1000) | 1.59 | 19.695 | 8.899 | 10.796 | 0.675 |
| Gefitinib (0 -25) | Trametinib (0 - 1000) | 0.402 | 10.254 | 4.513 | 5.741 | 0.55 |
| Buparlisib (0 -100) | Ibrutinib (0 - 1000) | 0.937 | 4.449 | 1.333 | 3.116 | 0.55 |
| Ibrutinib (0 -100) | Doxorubicin (0 - 100) | 0.84 | 9.822 | 1.734 | 8.088 | 0.525 |
| Vinorelbine (0 -1000) | Clofarabine (0 - 1000) | -1.143 | 66.463 | 21.399 | 45.064 | 0.5 |
| Clofarabine (0 -1000) | Omacetaxine (0 - 1000) | -4.266 | 20.336 | 0.585 | 19.751 | 0.375 |
| Dexamethasone (0 -1000) | Clofarabine (0 - 1000) | -16.368 | 0 | 0 | 0 | 0.35 |
| Dasatinib (0 -1000) | Ipatasertib (0 - 100) | 0.237 | 0.672 | 2.077 | -1.405 | 0.25 |
| Carboplatin (0 -1000) | Dexamethasone (0 - 1000) | -0.507 | 0 | 0 | 0 | 0.2 |
| Ipatasertib (0 -1000) | ASP3026 (0 - 1000) | -0.009 | 0 | 0 | 0 | 0.175 |
| Idarubicin (0 -100) | Ibrutinib (0 - 100) | -0.893 | 0.689 | 2.28 | -1.591 | 0.175 |
| Trametinib (0 -100) | Dasatinib (0 - 25) | -2.049 | 0 | 1.232 | -1.232 | 0.1 |

Synergy scores were calculated using ZIP [8] model (default option in SynToxProfiler).

To further illustrate the wide applicability of SynToxProfiler also in non-cancer combinatorial screens, we used a published dataset of 77 drug combinations tested as anti-viral agents where the drug combinations’ efficacy and toxicity were tested in Ebola-infected and non–virus-infected Huh7 liver cells, respectively. SynToxProfiler ranked established combinations (e.g. clomifene-sertraline and sertraline-toremifene) that inhibit EBOV fusion to cell surface as top hits for further study (Additional File 2). All the three drugs (clomifene, sertraline and toremifene) showed survival benefit in in-vivo murine Ebola virus infection model [18], indicating that SynToxProfiler prioritizes drug pairs with a strong potential to be rapidly advanced towards clinical settings and used as therapeutic interventions.

### Top hits selected by SynToxProfiler based on integrated scoring are synergistic drug pairs with higher selective efficacy

We compared the synergy and selective efficacy level of the top hits prioritized based on the STE score, synergy score and selective efficacy scores, using the 77 combinations in the Ebola dataset. The top combinations identified by STE scores had a notably higher selective efficacy as well as higher synergy (shown by arrow in Fig. 2A), indicating that STE score represents a proper balance between high selective efficacy and synergy. Additionally, we observed a marked overlap (65%) between the top-10% of analyzed combinations prioritized based on STE score and synergy score, as well as based on the STE score and selective efficacy score (50% overlap), as shown in Fig. 2B. In contrast, there was a smaller overlap (41%) between the top-10% hits selected based on selective efficacy and synergy scores. Further, a low Pearson correlation (r=0.22) between selective efficacy and synergy was observed. These results indicate that synergy and selective efficacy are independent drug combination components, which cannot be used alone to prioritize potent and less toxic synergistic drug combinations.

**Fig. 2.**
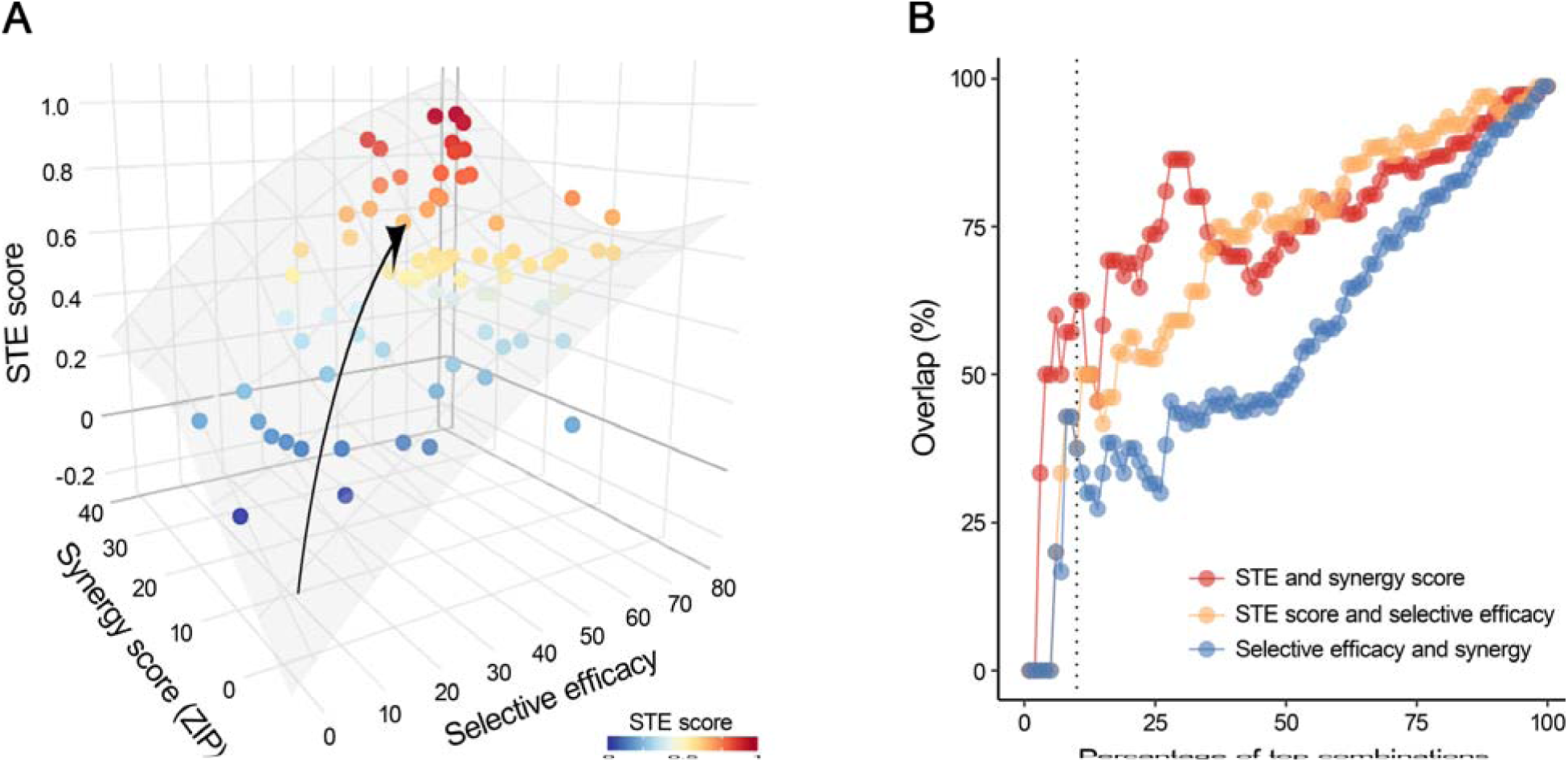
STE score considers both synergy and selective efficacy when prioritizing potent drug combinations. (A) 3D surface shows increase in STE score with increasing synergy and selective efficacy scores across 77 antiviral combinations measured in Huh7 liver cell line infected with Ebola virus (the arrow marks the gradient of the increase in STE score). The 3D surface is fitted by a generalized additive model with a tensor product smooth, implemented in mgcv R package. (B) Scatter plot showing the overlap in the top hits selected on the basis of different scores (the dotted vertical line denotes the overlap between the top 10% combinations selected based on any of the three scores).

A more detailed analysis revealed that SynToxProfiler ranks lower the toxic drug pairs despite their higher synergy (e.g. clomifene-colchicine and toremifene citrate-apilimod). For example, SyntoxProfiler ranked clomiphene citrate and sertraline HCl combination (STE=0.96) as the top hit (Fig.3), despite its lower synergy as compared to more synergistic toremifene citrate and apilimod pair (STE=0.86). This is due to a higher toxicity (13.30 vs 24.60) of latter, although both of the drug combinations have similar efficacy scores (70.88 vs 68.20). The lower ranking of combinations involving cilchicine and apilimod is in accordance with their observed extreme toxicity in the clinic [19,20]. This case study indicates that SynToxProfiler can identify safe top hits with high selective efficacy and synergy that have increased potential for clinical success, as compared to hits selected based on synergy alone.

**Fig. 3.**
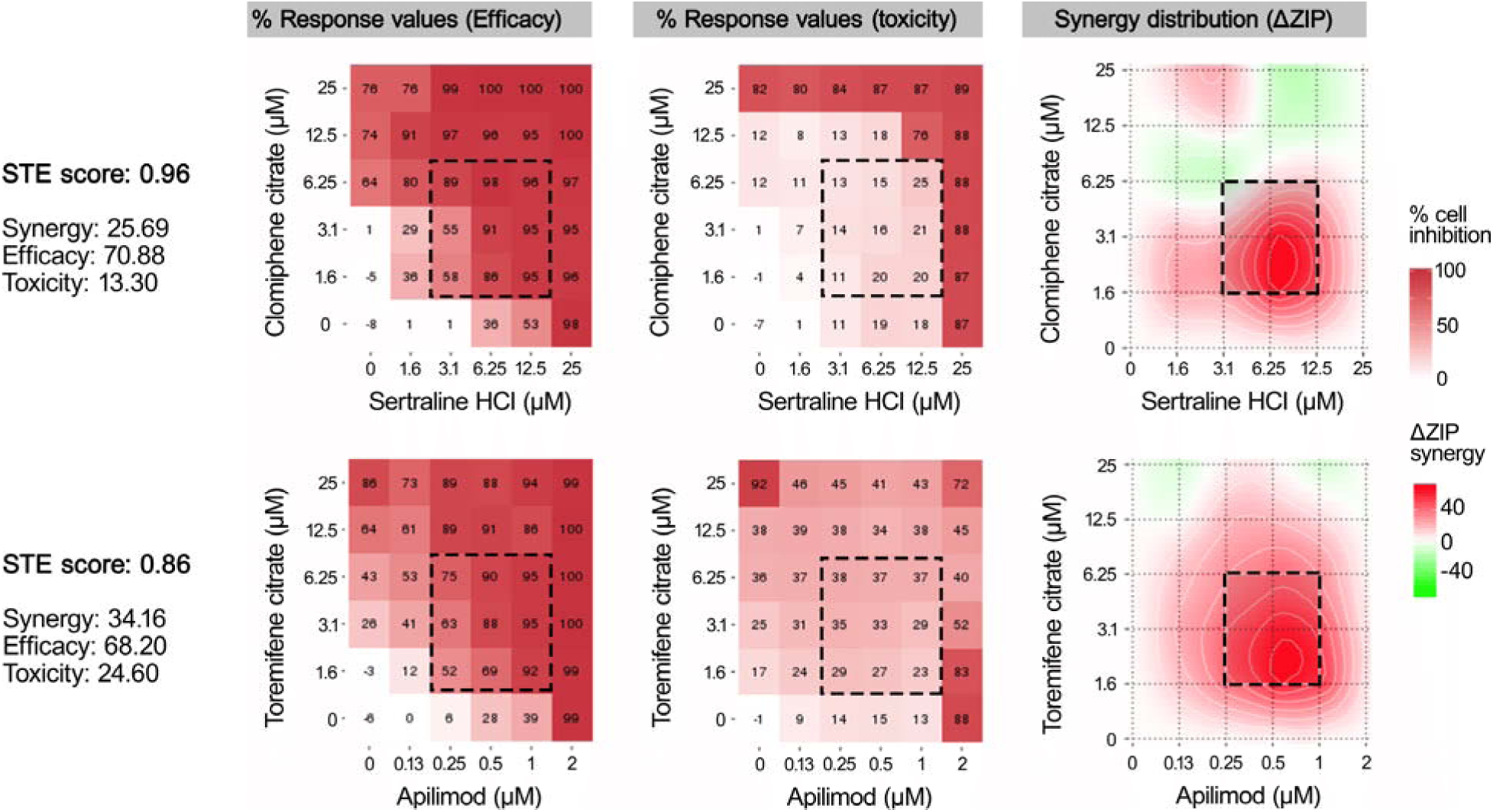
SynToxProfiler penalizes for toxicity of drug pairs while ranking top hits. (A) The efficacy, toxicity, and synergy matrices for the top drug pairs selected based on the highest STE score (clomiphene citrate and sertraline HCL, upper panel) and the highest synergy score (toremifene citrate and apilimod, lower panel). The synergy was calculated using the ZIP model implemented in SynergyFinder. The square with dotted line denotes the 3×3 concentration range with the most synergistic area in the dose-response matrix.

## Discussion and conclusions

The primary motivation for the use of synergistic drug combinations in the clinic is to achieve higher efficacy (by means of drug interaction) with reduced toxicity (by decreasing the drug doses). Therefore, the HTS screening aims to discover drug pairs that are more effective than the individual single drugs, and, at the same time, show less toxicity for the patients. Hence, the assessment of synergistic efficacy along with toxicity is critical for the selection of candidate drug pairs for further study, as there exists a fundamental trade-off between clinical efficacy and tolerable toxicity.

To the best of our knowledge, there are currently no methods to provide the global view in terms of synergy, efficacy and toxicity of drug pairs in an HTS setting. In this respect, SynToxProfiler offers an important advancement into the current practice for drug combination selection, as it provides an easy-to-use platform for in-vitro or ex-vivo assessment of the three critical aspects of drug combinations that are necessary for success in the clinics. Furthermore, SynToxProfiler facilitates the identification of therapeutic window range at which the drugs show highest synergy, high efficacy and lowest toxicity by visualization of the dose-response surfaces. Since, SynToxProfiler uses the normalized volume-based scoring for synergy, efficacy and toxicity levels (see methods and supplementary file), the SynToxProfiler framework can be easily utilized to prioritize synergistic drug combinations with high selective efficacy for multi-component (3 or more drugs) drug combination screening. Since limited number of tools and methodology are available to analyze and interpret either synergy, efficacy or toxicity of multi-component drug combinations, SynToxProfiler will be valuable resource for screening of such combinations.

In this work, we showed how SynToxProviler prioritized cytarabine-daunorubicin as the top drug pair out of 20 anticancer combinations for T-PLL case study (Table 1), and clomifene-sertraline for anti-viral case study (Additional file 2). The identification of clinically established drug pairs as top hit suggests that ranking based on all the three parameters can help to identify combinations that have more chance to success in the clinic. These effective and safe combinations would have been otherwise missed if combinations were selected merely based on their synergy scores.

In conclusion, we have developed SynToxProfiler, an interactive tool for top hit prioritization that ranks drug pairs based on their combined synergy, efficacy and toxicity profile, and which can be applied to any HTS drug combination screening project. We showed how this tool enable identification of clinically established drug pairs as top hits and many more drug pairs with a translational potential. We foresee SynToxProfiler will allow for more unbiased and systematic means to evaluate the pre-clinical potency of drug combinations toward safe and effective therapeutic applications.

## Authors’ contributions

AI and AKG developed and tested the integrated scoring. AI implemented the platform and AKG helped in designing and testing of the platform. ST performed the in-house drug combination screening in T-PLL case study. AI prepared figures for manuscript and finalized with AKG. TA helped in designing of the project and writing of the manuscript. AKG, TA, AI and AK conceptualized the study and wrote the manuscript. All authors have read and approved the final manuscript.

## Acknowledgements

We thank Prof. Satu Mustjoki for her valuable suggestions about the clinical use of SynToxProfiler, Prof. Krister Wennerberg for many discussions regarding synergy, toxicity and efficacy scoring approaches for drug combinations, and Andrea Cremaschi for valuable discussions and suggestions on volume-based combination scoring.

## Availability and requirement

The SynToxProfiler web-application is publicly available at https://syntoxprofiler.fimm.fi, together with drug combination example data, user instructions, and the source code. The source code is also available at https://github.com/IanevskiAleksandr/SynToxProfiler.

