## Supplementary Information for "SynToxProfiler: an approach for top drug combination selection based on integrated profiling of synergy, toxicity and efficacy"

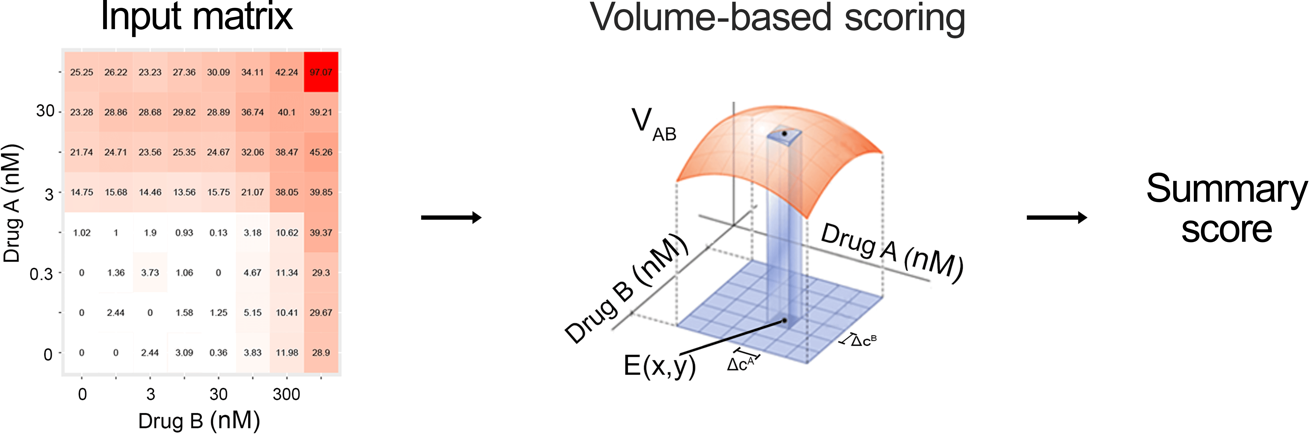

**Supplementary Fig. 1: Quantification of efficacy and toxicity in SynToxProfiler.** A schematic representation of calculation of *combination efficacy, synergy and toxicity* based on dose–response measurements on diseased cells or control cells. E(x,y) is the response at concentrations x and y of drugs A and B, respectively; Δc^A^ and Δc^B^ are the logarithmic increase in concentration of drug A and drug B between two consecutive measurements of the dose-response matrix.

**Supplementary Table S1: List of drugs used in the assay and their mechanism of action.**

| **Drug name** | **Targets** | **Mechanism** | **Approval status** |
| --- | --- | --- | --- |
| Ipatasertib | AKT inhibitor | Kinase inhibitor | Investigational |
| Daunorubicin | Topoisomerase II inhibitor | Chemotherapy | Approved |
| Doxorubicin | Topoisomerase II inhibitor | Chemotherapy | Approved |
| Omacetaxine | 80 S ribosome inhibitor | Chemotherapy | Approved |
| Navitoclax | Bcl-2/Bcl-xL inhibitor | Apoptotic modulator | Investigational |
| Idarubicin | Topoisomerase II inhibitor | Chemotherapy | Approved |
| S-63845 | MCL-1 inhibitor | Apoptotic modulator | Probe |
| Alpelisib | PI3Kalpha selective inhibitor | Kinase inhibitor | Investigational |
| Trametinib | MEK1/2 inhibitor | Kinase inhibitor | Approved |
| ASP3026 | ALK inhibitor | Kinase inhibitor | Investigational |
| Prexasertib | Chk1 inhibitor | Kinase inhibitor | Investigational |
| Dexamethasone | Glucocorticoid, immunomodulatory agent | Immunomodulatory | Approved |
| Clofarabine | Antimetabolite; Purine analog | Chemotherapy | Approved |
| Ibrutinib | Btk inhibitor | Kinase inhibitor | Approved |
| Dasatinib | Abl, Src, Kit, EphR Inhibitor | Kinase inhibitor | Approved |
| Cytarabine | Antimetabolite, interferes with DNA synthesis | Chemotherapy | Approved |
| Gefitinib | EGFR inhibitor | Kinase inhibitor | Approved |
| Carboplatin | Platinum-based antineoplastic agent | Chemotherapy | Approved |
| Vinorelbine | Mitotic inhibitor | Chemotherapy | Approved |
| Buparlisib | PI3K inhibitor | Kinase inhibitor | Investigational |
| Quizartinib | FLT3 inhibitor | Kinase inhibitor | Investigational |

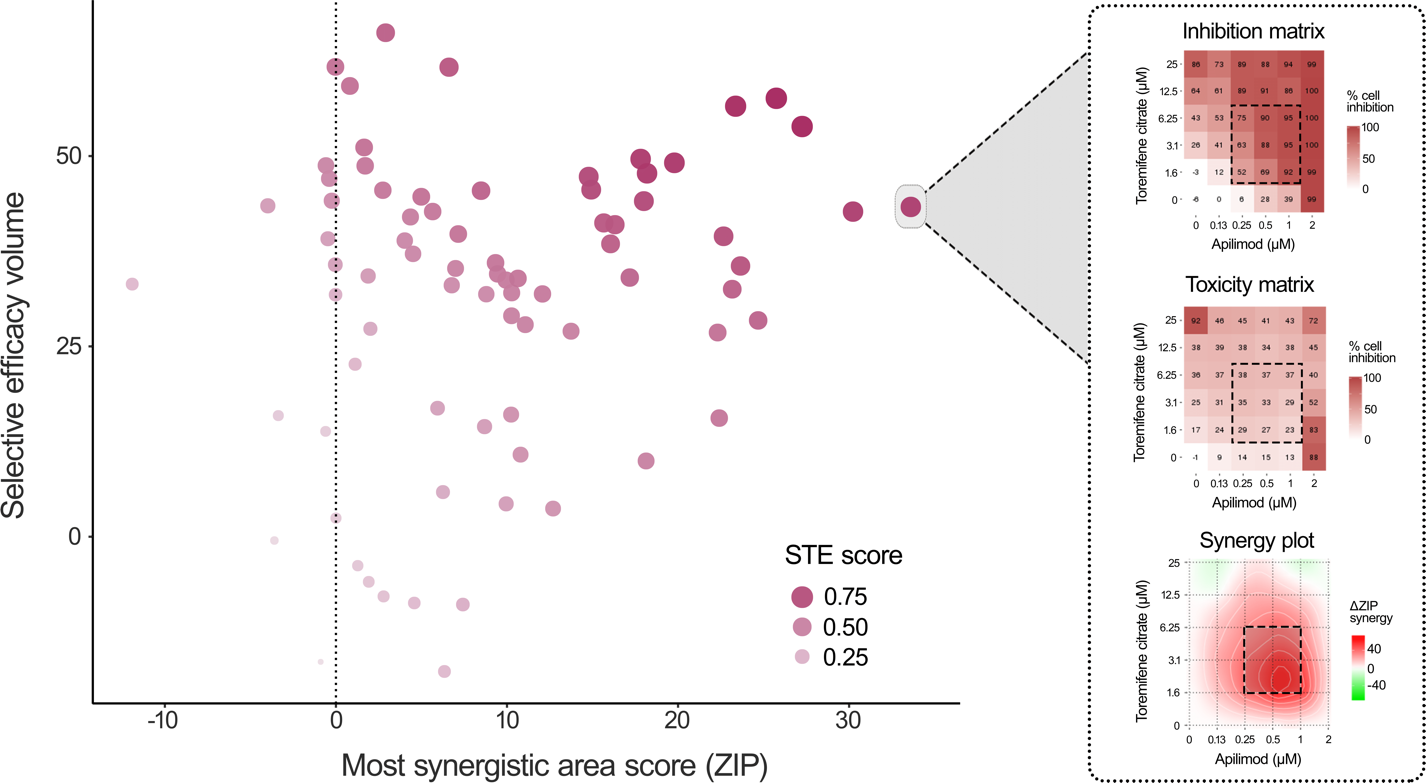

**Supplementary Fig. 2: Two-dimensional visualization in SynToxProfiler.** Scatter plot showing the distribution of a synergy score (x-axis) and selected efficacy score (y-axis) for 77 combinations tested in the Ebola infected and non–virus-infected Huh7 liver cells. Each drug combination is colored according to its STE score. Users can hover over the combinations to visualize their individual scores (e.g. STE score, or combination synergy, efficacy and toxicity scores), along with different dose-response matrices (synergy, toxicity, and efficacy), separately for each drug combination as shown here for the apilimod- toremifene citrate combination (right panel)

*Calculation of normalized volume score for multi-drug combinations*

For the case of multi-drug combinations (combination of 3 or more drug, the normalized volume under the multidimensional dose-response surface is calculated while quantifying combination efficacy and toxicity based on measurements on diseased and control cells, respectively. Synergy score is estimated based on measurements on diseased cells and expected combination responses are determined by one of the synergy models applicable for multi-drug combinations (e.g. Bliss or HSA). For example, for three-drug combination ABE of drugs A, B and E, the normalized volume V_ABE_ under the multidimensional dose-response surface is calculated as:

$$V_{\mathrm{ABE}}=\frac{\sum_{x=C_{\min}^{A}}^{C_{\max}^{A}} \sum_{y=C_{\min}^{B}}^{C_{\max}^{B}} \sum_{z=C_{\min}^{E}}^{C_{\max}^{E}} E(x,y,z)\Delta c^{A}\Delta c^{B}\Delta c^{E}}{\ln\left( C_{\max}^{A}/C_{\min}^{A} \right) ln(C_{\max}^{B}/C_{\min}^{B})ln(C_{\max}^{E}/C_{\min}^{E})} . Eq. (1)$$

Here, c^A^_min_ and c^A^_max_ are the minimum and maximum tested concentrations of drug A, respectively, and c^B^_min_, and c^B^_max_ are those of drug B and c^E^_min_, and c^E^_max_ are those of drug E; Δc^A^, Δc^B^ and Δc^E^ are the logarithmic increase in concentration of drugs A, B and E between two consecutive measurements of multidimensional dose-response matrix; and E(x, y, z) is the efficacy, toxicity or synergy levels at concentration x of drug A, concentration y of drug B and concentration z of drug E.
